# Tight junction component protein claudin-1 deficiency in retinal pigment epithelium leads to early and intermediate age-related macular degeneration phenotypes in mice

**DOI:** 10.1101/2023.12.07.570525

**Authors:** Ayaka Nakai, Deokho Lee, Yan Zhang, Chiho Shoda, Tetsu Imanishi, Heonuk Jeong, Shin-ichi Ikeda, Kazuno Negishi, Hiroyuki Nakashizuka, Satoru Yamagami, Akiharu Kubo, Toshihide Kurihara

## Abstract

The early and intermediate age-related macular degeneration (AMD) is characterized by the presence of drusen and pigmentary abnormalities in the retinal pigment epithelial (RPE) cells which form the outer blood retinal barrier (oBRB). Fluid leakage through the disrupted oBRB from the choroid to the neural retina has been implicated in the pathogenesis of AMD, however; the molecular mechanisms still remain unclear. The family of four transmembrane proteins, claudins are known to form tight junctions (TJs) in the oBRB. Nonetheless, there are few reports showing how they function in the oBRB *in vivo*. We found that claudin-1 is dominantly expressed in TJs of the mouse RPE. To investigate the role of claudin-1 in the RPE, we generated RPE-specific *Cldn1* conditional knockout mice (Best1-Cre^+/-^ *Cldn1*^flox/flox^ mice: *Cldn1* cKO mice). Deficiency of *Cldn1* led to age-related lipid deposits such as subretinal drusenoid deposits (SDD), increased lipid droplets in the RPE, basal lamellar deposits (BlamD) and membranous debris in the Bruch’s membrane. In addition, pigmentary abnormalities such as RPE hypertrophy, multilayered-RPE cells, and ectopic pigment granules outside the RPE were observed in *Cldn1* cKO mice. Our study provides new insights into the possible association of the TJ protein claudin-1 with lipid metabolism and cellular ageing in the RPE contributing to the early onset of AMD.

## Introduction

The retinal pigment epithelium (RPE) is a single layer epithelium located between the neural retina and choroid. Photoreceptor cells consisting the outermost layer of the neural retina are in contact with RPE cells and nourished by choroidal vessels (Runkle and Antonetti, 2011). The RPE cells has three main functions, a pump function that constantly transports fluid within the retina to the choroid, control of paracellular pathway by forming the outer retinal blood-retinal barrier (oBRB), and phagocytosis and degradation of photoreceptor outer segments. The Bruch’s membrane exists between the RPE and the choroid and consists of 5 layers as RPE basal lamina, inner collagenous layer, elastic layer, outer collagenous layer, and choriocapillaris basal lamina from the retinal side. RPE cells adhere closely to adjacent cells and form oBRB (Cunha-Vaz, 1976; Kaur, Foulds and Ling, 2008; Runkle and Antonetti, 2011). The oBRB selectively transfers fluid, nutrients, oxygen and metabolites between choriocapillaris, the most inner layer of the choroid, and neural retina bilaterally (Kaur, Foulds and Ling, 2008; Naylor et al., 2020).

The oBRB is mainly composed of tight junctions (TJs) with specific protein complexes. TJs contain a complex arrangement of transmembrane proteins and cytoplasmic scaffolding proteins (Runkle and Antonetti, 2011; Naylor et al., 2020). Deletion of the transmembrane protein families of claudin and occludin, and the scaffolding protein zonula occludens-1 (ZO-1), was suggested to associate with increases in permeability in oBRB (Xu and Le, 2011; Peng et al., 2016; Zhang et al., 2019). It has been suggested that oBRB leakage is involved with retinal edema and visual dysfunction in retinal diseases such as AMD (Bernardes et al., 2011; Joussen et al., 2021), central serous chorioretinopathy (CSC) (Eandi et al., 2005; Naylor et al., 2020), and diabetic retinopathy (DR) (Xu and Le, 2011; Omri et al., 2013; Joussen et al., 2021).

Claudins are four-transmembrane proteins (Furuse et al., 1998) and 27 of family members have been identified (Wu et al., 2006; Mineta et al., 2011; Günzel and Yu, 2013). Claudin-1 is a transmembrane protein responsible for maintaining the epithelial barrier mechanism at tight junctions in epithelial cells. There are no reports regarding the relationship between claudin-1 in RPE and aging. Claudin-19 is known to be involved with the maintenance of oBRB proper permeability, destiny of the RPE, and gene expressions for the RPE homeostasis in the human retina (Wang et al., 2019; Liu et al., 2021). However, roles and mechanisms of claudins in the RPE are still poorly understood.

The early and intermediate AMD are clinically diagnosed by the presence of drusen and/or RPE pigmentary abnormalities without symptoms (Ferris et al., 2013; Mitchell et al., 2018). AMD eventually progresses to late AMD, classified as neovascular AMD and geographic atrophy (GA), which can cause visual impairment. Drusen is a sub-RPE accumulation of lipids, apolipoproteins, sugar, and inflammatory substances such as clusterin, complement C3, and amyloid-β (Holz et al., 1994; Haimovici et al., 2001; Sakaguchi et al., 2002; Curcio, 2018). The advances in imaging and pathological research have identified various types of extracellular deposits such as basal lamellar deposits (BlamD: a thickened basement membrane of the RPE), soft drusen and basal linear deposits (BlinD: accumulated between the RPE basal lamina and inner collagenous layer, and subretinal drusenoid deposits (SDD: located between the RPE and photoreceptors) (van der Schaft et al., 1993; Curcio, 2018). SDD is cholesterol-rich debris in the subretinal spaces containing lipid oil droplet. Morphologically, it formed irregular oval inclusions less than 1 μm in diameter, superficially resembling a condensate of outer segment-like material but lacking internal structure similar to photoreceptor outer segment (OS) discs (Curcio et al., 2005, 2013). Pigmentary abnormalities is divided into hypopigmentation and hyperpigmentation by fundus findings. RPE cells from AMD donor eyes show morphological abnormalities of the RPE cells, such as enlargement (Watzka et al., 1993; Gambril et al., 2019; Ortolan et al., 2022) and diversification (Gambril et al., 2019; Ortolan et al., 2022).

It is important to examine extracellular lipid deposits such as BlamD, soft drusen, BlinD, and SDD, as well as pigmentary abnormalities for understanding the pathogenesis and progression of AMD. In this study, we investigated the expression of the tight junction (TJ) transmembrane protein claudins in the mouse RPE. Moreover, we revealed the function by generating RPE-specific conditional knockout mice.

## Materials & Methods

### Animals

Mice were maintained in a 12-hour light-dark cycle and with ad libitum access to food and water. All wild type C57BL/6J mice were purchased from CLEA Japan Inc (Tokyo, Japan). *Cldn1*^flox/flox^ mice were obtained as described in Figure 1. To eliminate to the neomycin resistance array inserted between FLP recombinase target (FRT) cassette, *Cldn1*^flox/flox^ mice (accession number: CDB0803K, preserved in Riken, Japan) were mated with *C57BL/6-Tg(CAG-flpe)36Ito/ItoRbrc* mice (accession number: RBRC01834, provided from Riken, Japan) with the *pCAGGS* expression vector provided by Dr. Jun-ichi Miyazaki ( Niwa, H. et al., 1991). Details about *Cldn1*^flox/flox^ mice is in the previous paper (Atsugi, T. et al., 2020). The detailed information on the mouse from Riken is described in the following URL: https://large.riken.jp/distribution/mutant-list.html

**Figure 1.**
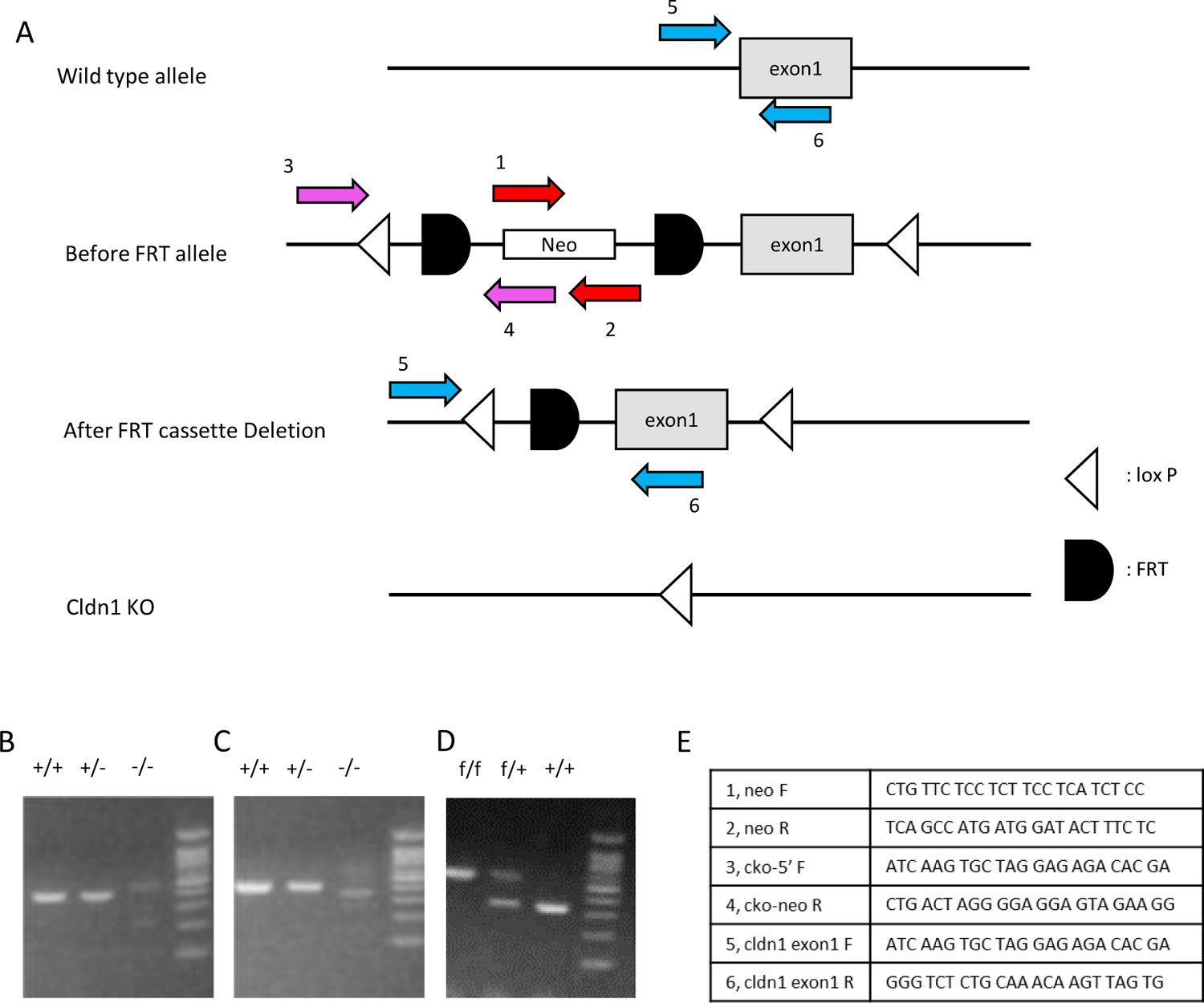
Generation of RPE specific claudin-1deficient mice. (A) Schematic representation of the transgene. FLP remove FLP recombinase target (FRT) cassette from before FRT allele. The red, pink and blue arrows indicate locations of attaching primers for genotyping. (B,C) Genotype analyses of the DNA extracted from *cldn1*^flox/flox^ mice before deleting FRT cassette, homozygous (+/+), heterozygous (+/-) and wild-type (-/-) mice for the mutant claudin-1 gene allele. (D) Genotype analyses of the DNA extracted from *cldn1*^flox/flox^ mice after FRT cassette dilation, homozygous (f/f), heterozygous (f/+) and wild-type (+/+) mice for the mutant claudin-1 gene allele. (E) The list of primers used for genotyping in the process of creating *Cldn1*^flox/flox^ mice.

*Best1-Cre* mice (Iacovelli et al., 2011) were purchased from the Jackson Laboratory (ME, USA). Mice with homozygous conditional inactivation of *Cldn1* gene in RPE cells were generated by breeding *Best1-Cre* transgenic mice with mice bearing a loxP-flanked *Cldn1* allele after FRT cassette deletion to create the first generation of *Best1-Cre* positive, *Cldn1* heterozygous mice (*Cldn1*^flox/–^) on a C57BL/6 background. The first generation of *Best1-Cre* positive *Cldn1* heterozygous mice (*Best1-Cre*^+/-^ *Cldn1* ^flox/–^) were subsequently crossed with *Cldn1* homozygous floxed mice (*Cldn1*^flox/flox^) to obtain *Best1-Cre*^+/-^ *Cldn1*^flox/flox^ mice (*Cldn1* cKO mice). *Best1-Cre*^-/-^ *Cldn1*^flox/flox^ mice were used as control mice in this study. All procedures were approved by the Ethics Committee on Animal Research of the Keio University School of Medicine (accession number: A2022-069) adhered to the Association for Research in Vision and Ophthalmology Statement for the Use of Animals in Ophthalmic and Vision Research, the Institutional Guidelines on Animal Experimentation at Keio University, and the Animal Research: Reporting of *In Vivo* Experimental guidelines for the use of animals in research. Genotyping was performed by PCR analysis with mouse tails. GoTaq DNA polymerase and reagents (Promega, USA) were used to amplify the products. The PCR conditions were as follows: initial denaturation (94◦C for 2 min), followed by 30 cycles of denaturation (94◦C for 30 sec), annealing (59◦C for 30 sec) and extension (72◦C for 1 min) and extension (72◦C for 5 min). The PCR products were separated on a 2.0% agarose gel. The primers used were in Figure 1 and as follows: *Best1-Cre* (forward: ATGCGCCCAAGAAGAAGAGG AAGGTCTCC, reverse: TGGCCCAAATGTTGCTGGATAGTTTTTA); *B6-Tg(CAG-FLPe)36* (*5CAG:*CCTACAGCTCCTGGGCAACGTGC, *Flp/Int.R*: CTGCTTCTTCCGATGATTCG).

### Swept source-optical coherence tomography (SS-OCT) and scanning laser ophthalmoscope (SLO) for image acquisition and measurement of retinal and choroidal thickness

Swept source-optical coherence tomography (SS-OCT) examinations using multiline OCT (Xephilio OCT-S1, Canon, Japan) were performed in 4, 8, and 16-week-old control and *Cldn1* cKO mice. Tropicamide and phenylephrine hydrochloride solution (Mydrin-P ophthalmic solution, Santen Pharmaceutical) was applied to the mouse eye 5 min before the measurement. The scanning was performed after the general anesthesia was induced by a combination of midazolam (Sandoz K.K.), medetomidine (Domitor, Orion Corporation), and butorphanol tartrate (Meiji Seika Pharma Co., Ltd.) which is called “MMB”. The images containing the center of the optic nerve head were used for the analysis. The retinal and choroidal thickness was measured by the manual assessment of the B-scan images. The distance between the ganglion cell layer (GCL) to the RPE was calculated as retinal thickness, and RPE-Bruch’s membrane-choroid complex was calculated as choroidal thickness. Retinal and choroidal thickness was assessed at distances of 0.8, 1.6, 3.2, 6.4 and 9.6 mm nasally and temporally from the optic nerve head. SLO examinations performed using the Xephilio OCT-S1machine (Canon, Japan) with the same anesthesia method above.

### Hematoxylin and eosin staining for retina/choroid/sclera cross-sectional histology and its immunohistochemistry and oil red O staining

Mice were deeply anesthetized using isoflurane inhalation and euthanized. The eyes were immediately enucleated, and the extrinsic ocular muscles were stripped off. The eyeballs were embedded in OCT compound (Sakura Tissue-Tek, Tokyo, Japan), and were rapidly frozen in liquid nitrogen. 8μm cyro samples were sectioned by microtome cryostat (CM3050S; Leica, Wetzlar, Germany). Sectioned samples were fixed for 5 min at room temperature in 4% paraformaldehyde and washed three times with phosphate-buffered saline (PBS). The section was assessed by hematoxylin and eosin staining, immunostaining and oil red O staining.

For hematoxylin and eosin staining, the samples were immersed in hematoxylin (MUTO PURE CHEMICALS CO.,LTD., Tokyo, Japan) for 1 min and eosin (MUTO PURE CHEMICALS CO.,LTD., Tokyo, Japan) for 10 sec followed by several washing.

For immunostaining, samples were permeabilized with PBS containing 0.2% Triton X-100 and 0.1% bovine serum albumin (BSA) for 40 min at room temperature. The samples were washed three times with PBS and blocked with 10% normal donkey serum for 1 h at room temperature. Primary antibodies were applied in 10% donkey serum at 4◦C overnight. After washing three times in PBS (at 5 min interval), samples were incubated with secondary antibodies for 1 h at room temperature. DAPI (DAPI-Fluoromount-G®, SouthernBiotech, Birmingham, AL, USA) was used for nuclear staining. The following antibodies were used in the current study: anti-claudin-1 (1:50, Cat #51-9000; Thermo Fisher Scientific, Waltham, MA, USA), anti-ZO-1 (1:100, Cat #14-9776-8; Thermo Fisher Scientific, Waltham, MA, USA), anti-RPE65 (1:100, Cat # NB100-355; Novus Biologicals, Centennial CO, CO, USA), and anti-F4/80 (1:100, Cat #123117123 101; BioLegend, San Diego, CA, USA) and appropriate Alexa Fluor-conjugated secondary antibodies were used (Thermo Fisher Scientific). Images were obtained using a confocal microscope (LSM710 or 980; Carl Zeiss; Oberkochen, Germany).

For oil red O staining, melanin of part of samples were breached before oil red O staining using Melanin Bleach Kit (Cat # 24883 Polysciences, Inc., Warrington, PA, USA). After washing in distilled water, samples were stained as followed by the general manufacture’s introduction. The samples were soaked in 60% isopropanol for 2 min and immersed in 60% oil red O regent (MUTO PURE CHEMICALS CO.,LTD., Tokyo, Japan) diluted by distilled water for 5 min. After fractionated with 60% isopropanol for 2 min, the samples were washed with distilled water and counterstained using hematoxylin for 1 min to visualize nuclei. Then, samples were rinsed with running tap water for 3 min and covered with a coverslip using glycerol.

### Flat-mount immunohistochemistry for RPE morphological analysis

For flat-mount immunostaining of the RPE flat-mount, mice were deeply anesthetized using isoflurane inhalation and euthanized. The eyes were removed and pre-fixed for 10 min on ice in 4% paraformaldehyde. Three incisions were made in the cornea, and the cornea, iris and lens were removed completely. From the eyecup, the sensory retina was removed using micro tweezers. The remaining RPE-choroid-sclera complex was cut open at from 5 to 6 leaves. After fixing with 4% paraformaldehyde for 1 h, the sample was washed with PBS and blocked with PBS containing 0.5% Triton X-100 and 1% BSA at 4◦C overnight. Primary antibodies were applied in PBS containing 1% BSA at 4◦C for three days, washed three times in PBS (20 min per wash), and followed by incubation with secondary antibodies at 4◦C overnight. Finally, the RPE flat-mount was washed three times with PBS (20 min per wash) and mounted in Fluoro-Gel (Electron Microscopy Sciences) solution. The antibodies used in this procedure were as same as written in the section immunostaining above. Images were obtained using a confocal microscope (LSM710 and LSM980; Carl Zeiss; Oberkochen, Germany).

For RPE morphological analysis, the samples were stained by anti-ZO-1 antibody after the RPE flat-mount procedure. The Images were observed with a fluorescence microscope (BZ-9000, KEYENCE, Osaka, Japan). A visual field (VF) was defined as a 960×720pixel photograph taken at the same angle of view and magnification. The number of cells in one VF and the area of each cell in VF were obtained using ImageJ (Open-source tools: https://imagej.nih.gov/ij/download.html). The average cell area for each VF and the coefficient of variation (CV) were calculated. The CV calculated as the standard deviation of cell area divided by the average cell area in each VF.

### *in vivo* permeability assay

After general anesthesia by MMB, mice were perfused with 0.2mg 70,000MW FITC-dextran per body weight (g) via the right retro-orbital injections. The injection method was followed as in the previous literatures (Li et al., 2011; Yardeni et al., 2011). After 5 min reflux, the mice were euthanized. Then, the left eye was enucleated and the extrinsic ocular muscles were stripped off. The eyeballs were embedded in OCT compound and frozen in liquid nitrogen. Sample sections were prepared in the same manner as for the cross-section above. After fixation with 4% paraformaldehyde for 5 min, the samples were washed two times in PBS (3 min per wash) and embedded in mounting medium containing DAPI. The Images were observed with a fluorescence microscope (BZ-9000, KEYENCE, Osaka, Japan).

### RPE isolation and Quantitative PCR

Primary mouse RPE cells were prepared following a previous report (Fernandez-Godino, Garland and Pierce, 2016). The 8-week-old and 12-week-old wild mouse eyes were removed and washed in HBSS without Ca^2+^ or Mg^2+^ using 10 mM HEPES. Three incisions were made in the cornea, and the iris and lens were completely removed. The eyes were placed in HEPES-free HBSS buffer and incubated in hyaluronidase solution at 37◦C for 45 min in a 5% carbon dioxide to detach the sensory retina from the RPE. Each eye was placed in a new well with 1.5mL of cold HBSS HEPES buffer per well and incubated on ice for 30 min. After washing, each eye was placed in a 12-well plate containing fresh HBSS HEPES buffer. After the sensory retina was pulled away, each eye cup was transferred to a different 12-well plate containing fresh trypsin-EDTA (1.5 mL) per well and incubated the eyecups at 37◦C for 45min in a 5% carbon dioxide incubator. Each eye cup was held by the optic nerve and shaken face down into a 12-well plate containing 1.5mL of 20% FBS in HBSS HEPES buffer until a complete detachment of the RPE sheets were achieved. RPE cells from the six eyes of three 8-week-old mice (Figure 2A) and four eyes of two 12-week-old mice (Supplementary Figure S4) were used for each sample. The RPE cells were dissolved in TRI reagent (MRC Global, Cincinnati, OH, USA) and RNA extraction was performed using EconoSpin® columns (GeneDesign, Osaka, Japan). The columns were washed with RPE and RWT buffers (Qiagen, Hilden, Germany). Reverse transcription from the extracted RNA was performed using the ReverTra Ace™ qPCR RT Master Mix with gDNA Remover (Toyobo Co., Ltd. Osaka, Japan). Quantitative PCR for analyzing gene expression was performed using THUNDERBIRD® SYBR® qPCR Mix (Toyobo Co., Ltd. Osaka, Japan) with the QuantStudio 5 (Life Technologies, Carlsbad, CA, USA). The primer sequences used in this study are listed in Supplementary Table S1. The fold change between the levels of different transcripts was calculated by the ΔΔCT method.

**Figure 2.**
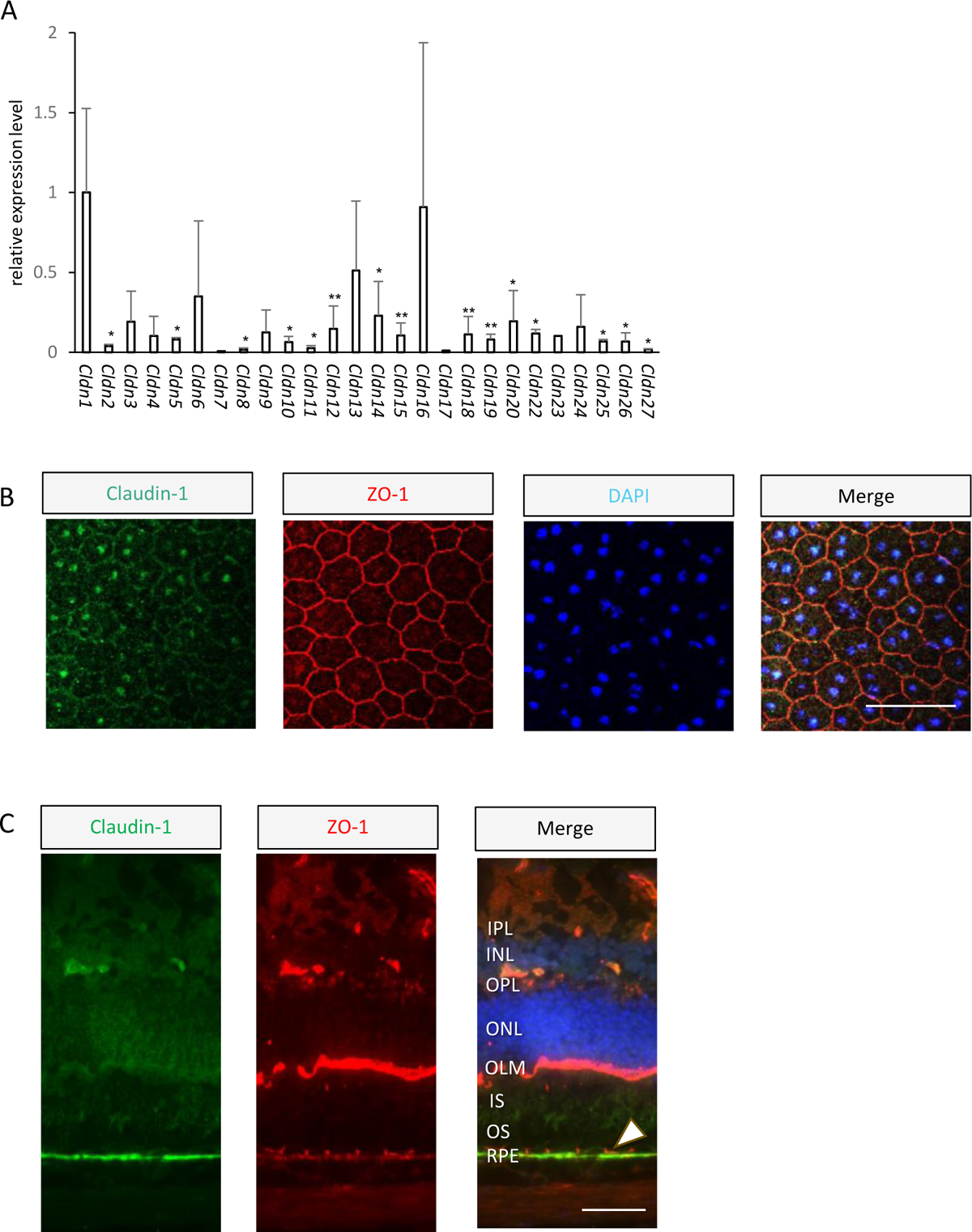
Claudin-1 expression in the outer retinal blood burrier (oBRB) of wild type mice (C57BL/6) (A) The mRNA expression levels of claudins in RPE isolated from 8-week-old Wild type mice (± SD, n=3, 1sample 6eyes). Student t test, vs *Cldn1*, *P <0.05,**P <0.01. (B) RPE flat-mounts sample from 8-week-old representative mouse and (C) cryo section from 1-yr-old representative mouse were stained with claudin-1 (green), ZO-1 (red) and DAPI (blue) and obtained by confocal microscopy. Claudin-1 was collocated with ZO-1 in the oBRB (head arrow). Scale bar, 50 μm. WT, Wild type; RPE, retinal pigment epithelium; OS, outer segments; IS, inner segments; OLM, outer limiting membrane; ONL, outer nuclear layer; OPL, outer plexiform layer; INL, inner nuclear layer; IPL, inner plexiform layer.

**Figure 3.**
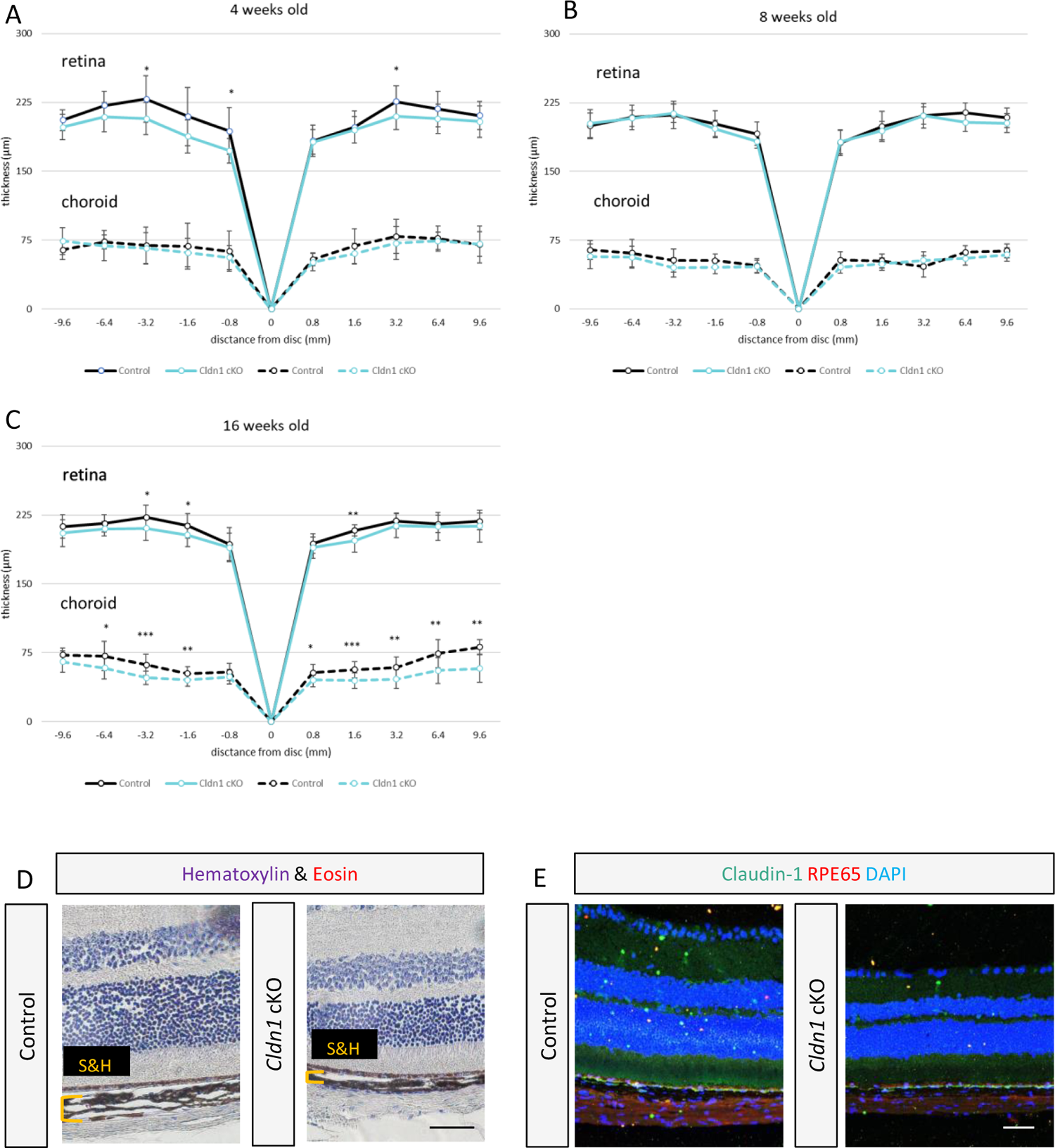
The retinal and choroidal thickness changed in *Cldn1* cKO mice. (A-C) The measurements of retinal and choroidal thickness in distance from the optic nerve (–0.96 mm: nasal, +0.96 mm: temporal). The thickness were obtained by measuring optical coherence tomography (OCT) images in 4, 8 and 16-week-old control and *Cldn1* cKO mice. (± SD, n=8, 4-week-old control; n=16, 4-week-old cKO mice; n=9, 8-week-old control; n=11, 8-weeks-old cKO mice; n=13, 16-week-old control; n=19,16-week-old cKO mice) In some images, the outermost part was not measurable. Student t test, *P <0.05,**P <0.01, ***P <0.001. (D,E) Representative images of retina/choroid/sclera cross sections. Scale bar, 50 μm. (D) Hematoxylin and eosin staining of 10-week-old mice. Sattler’s and Haller’s layer (bracket) were thinner in *Cldn1* cKO mice choroid. (E) immunohistochemistry of 20-week-old mice stained with claudin-1 (green), RPE65 (red) and DAPI (blue).

### Transmission electron microscopy

The mice were deeply anesthetized by MMB and perfused with saline mixed 2.5% glutaraldehyde. The eyes were immediately enucleated and the corneas were removed and fixed in 2.5% glutaraldehyde in PBS (0.1 M, pH 7.3) overnight. The lens and extrinsic ocular muscles were removed, and the posterior segment (the retina, choroid, and sclera) was halved with knife to contain the perioptic nerve portion at the apex and ciliary body at the base. The halved samples were submitted for electron microscopic sectioning. The tissue was fixed in 1% osmium tetroxide for 1 h, rinsed in PBS, dehydrated in EtOH and then embedded in Spurr’s resin. Thick (0.7–1.0 μm) and ultrathin sections (0.1 μm) were cut on a microtome (Porter Blum MT-2). Thick sections were stained with toluidine blue and examined by light microscopy. Ultrathin sections were stained with 4% uranyl acetate and lead citrate and then examined with a CX-100 transmission electron microscope (JEOL, Tokyo, Japan).

### Statistical analysis

A two-tailed Student’s *t*-test (Excel, Microsoft, WA, USA) was used to compare the mean variables of the two groups. All data were presented as the means ± standard deviation. Statistical significance was set at p < 0.05.

## Results

### *Cldn1* was dominantly expressed in tight junctions (TJs) of the mouse RPE

First, we evaluated which number of the claudin genes might be the most highly expressed among 27 claudin family members in the mouse RPE. *Cldn1* was primarily expressed in RPE cells isolated from wild type (WT) mice (Figure2 A, Supplementary Figure S4). In addition, *Cldn16* was also relatively highly expressed. ZO-1-positive polygonal mesh structure was also seen in our samples (Figure2 B). Claudin-1 was collocated with ZO-1 in the RPE of the flat mount and the vertical retina section (Figure2 B,C). We found that claudin-1 is expressed in the TJs of the RPE, forming the oBRB.

### Claudin-1 deficiency in mouse RPE induced age-related morphological abnormalities in RPE promoting thinning of the retina and choroid

To explore the function of claudin-1 *in vivo*, we generated RPE-specific *Cldn1* conditional knockout mice (*Cldn1* cKO mice). We found that *Cldn1* cKO mice showed thinner retina and choroid *in vivo* live image by optical coherence tomography (OCT) (Figure3 A-C). Retinal thickness was significantly thinner in 4- and 16-week-old *Cldn1* cKO mice, and choroidal thickness was thinner in 16-week-old *Cldn1* cKO mice (Figure3 A,C). especially middle and large vascular layers so called Sattler’s and Haller’s layer compared to the control in histological section (Figure3 D,E). It is known that AMD patients and aged mice have morphological changes in RPE such as the large cell area and a variability of cell shape (Chen et al., 2016; Gambril et al., 2019; Ortolan et al., 2022). *Cldn1* cKO mice showed regional morphological changes in the RPE (Figure 4 A,B). Specifically, large size cells, round misshapen small cells, and large multangular cells were observed in *Cldn1* cKO mice (Figure 4B). 8-week-old *Cldn1* cKO RPE showed significantly smaller cell numbers (p<0.001) (Figure 4C) and larger cell area in central area (p<0.001) (Figure 4D). The diversity of individual cell area was larger in equatorial (p<0.01) and peripheral areas (p<0.05) compared to the control (Figure 4E). Similar outcomes were also observed in 20-week-old *Cldn1* cKO mice (figure 4F-H). The difference in CV between Control and Cldn1 cKO mice in 20-week-old mice was larger than that of in 8-week-old mice, with an average of 0.13 vs. 0.03 in three areas (central, equatorial and peripheral areas). Taken together, claudin-1 deficiency may accelerate age-related morphological changes in the RPE, promoting thinning of the retina and choroid.

**Figure 4.**
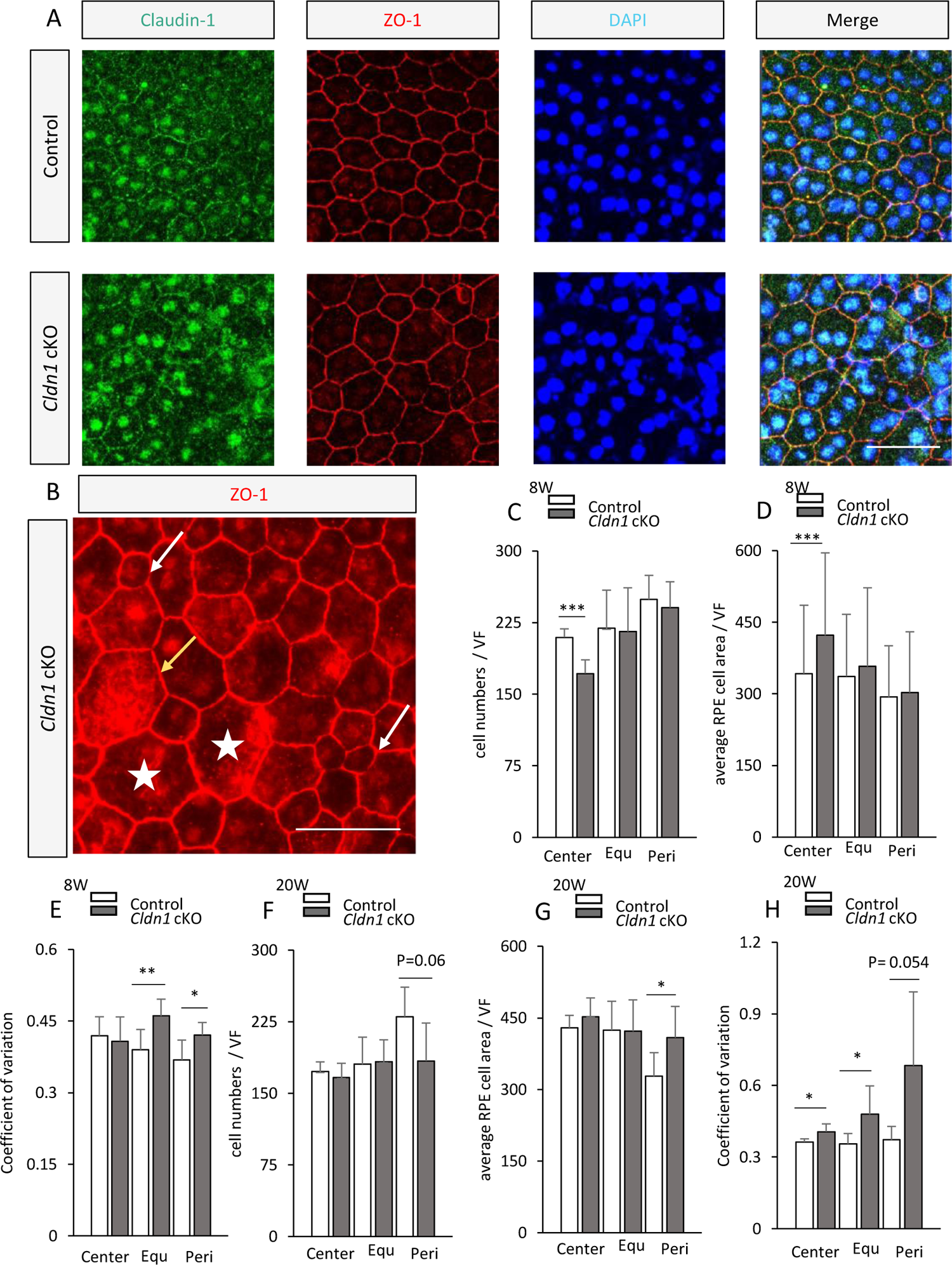
Age-dependent RPE surface morphologic abnormality in *Cldn1* cKO mice. (A) RPE flat-mounts from 8-week-old control and *Cldn1* cKO mouse were stained with claudin-1 (green), ZO-1 (red) and DAPI (blue). (B) 8-week-old *Cldn1* cKO mouse high-magnification flat mount image showed a giant RPE cell (yellow arrow), small less angular cells (white arrow) and octagon cells (star). (C-H) The RPE morphological analysis was performed. Quantification of (C,F) cell numbers, (D,G) cell area and (E,H) coefficient of variation per the visual field (VF) was performed using ImageJ from ZO-1 stained RPE flat-mounts image (± SD, n=5-7 VF per group). Student t test, *P <0.05,**P <0.01, ***P <0.001. Equ, equatorial; Peri, peripheral. (A,B) Scale bar, 50 μm. VF: visual field.

### *Cldn1* cKO mice showed hyperpigmentary abnormalities with RPE hypertrophy, multiple layered-RPE and ectopic pigment granules

Drusen and pigmentary abnormalities could lead to AMD progression (Wang et al., 2003; Ferris et al., 2013). Pigmentary abnormalities were divided into hyperpigmentation and hypopigmentation. RPE hyperpigmentation may result in RPE cell hypertrophy, multiple layered-RPE, or ectopic pigment granules. We observed patchy RPE hypertrophy in adult *Cldn1* cKO mice (Figure 5A,B). In transmission electron microscopy (TEM) images, RPE cells were aligned in a row on the Bruch’s membrane in the control mice (Figure 5C). In contrast, *Cldn1* cKO mice showed multiple layered RPE and accumulation of lipofuscin–like vacuoles within the RPE (Figure 5D). RPE organelles were deviated from RPE cells and ectopic pigment granules were observed in the extracellular space (Figure 5E).

**Figure 5.**
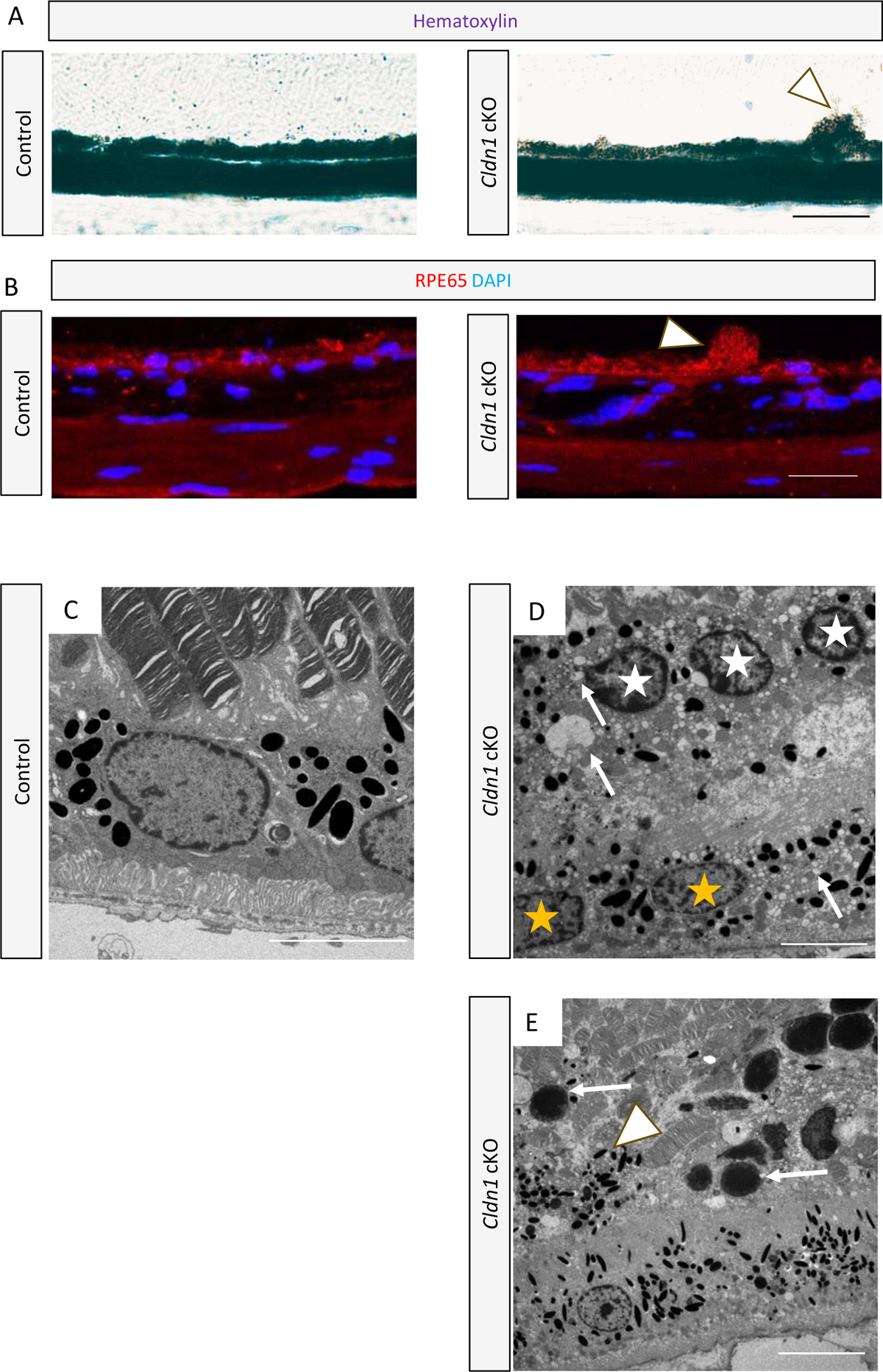
Hyperpigmentary abnormality observed in *Cldn1* cKO mice RPE. (A,B) Patchy RPE hypertrophy (head arrows) were observed in adult Cldn1 cKO mice. (A) Hematoxylin and oil red o staining of control and *Cldn1* cKO 20-week-old mouse. Scale bar, 20 μm. (B) 20-week-old control and cKO mouse were stained with RPE65 (red) and DAPI (blue). Scale bar, 20 μm. (C-E) TEM images of RPE in control and *Cldn1* cKO 16-week-old mice. (n = 6 eyes per group) (C) Control. The RPE cells maintain a monolayer structure, and the organelle exists only within the RPE. Scale bar, 5 μm. (D) The RPE shows facing Bruch’s membrane (yellow stars). Besides the RPE multi-layer (white stars) is observed in *Cldn1* cKO mice. Various sizes vacuoles (arrows) are present within the RPE. Scale bar, 5 μm. (E) RPE organelle including melanin (head arrow) is observed extending out of the monolayer RPE structure in *Cldn1* cKO mouse. Photoreceptor nuclei (arrows) are observed ectopically. Scale bar, 10 μm.

### *Cldn1* cKO mice showed age-dependent accumulation of lipids, lipofuscin–like vacuoles in the RPE; basal laminar deposits; and membranous debris in the Bruch’s membrane

Abnormal accumulation of lipids was seen within RPE in *Cldn1* cKO mice (Figure5 D). Therefore, analyses on lipids around the RPE and Bruch’s membrane were focused. We found that lipid droplets were detected in the RPE of *Cldn1* cKO mice (Figure 6A). BlamD and type 1 macular neovascularization, a type of the neovascular AMD in which neovascular vasculature are located under the RPE, showed two highly reflective layers on the OCT images; RPE and beneath the RPE (Sato et al., 2007; Sura et al., 2020). The two reflective bands are named as the “double-layer sign”. SLO images showed a circular hyperreflective region with irregular edges (Figure 6B), and the OCT image of the same area showed a double-layer sign (Figure 6B). Compared to control mice (Figure 6C), TEM revealed thickened basal infoldings (BI) and BlamD (Figure 6D,E). The TEM outcomes may suggest that the OCT image was due to BlamD, an abnormal lipid accumulation in the RPE basal lamina of *Cldn1* cKO mice. Membranous debris in outer collagenous layer of the Bruch’s membrane was also often observed in *Cldn1* cKO mice (Figure 6F), whereas the Bruch’s membrane in control mice exhibited uniformed thickness (Figure 6C). The membranous debris was present in 4 of 6 *Cldn1* cKO mice eyes and BlamD in 6 of 6 eyes. In *Cldn1* cKO mice, lipid deposition was present in the RPE, sub-RPE, and Bruch’s membrane. BlamD was also detected by a live OCT image.

**Figure 6.**
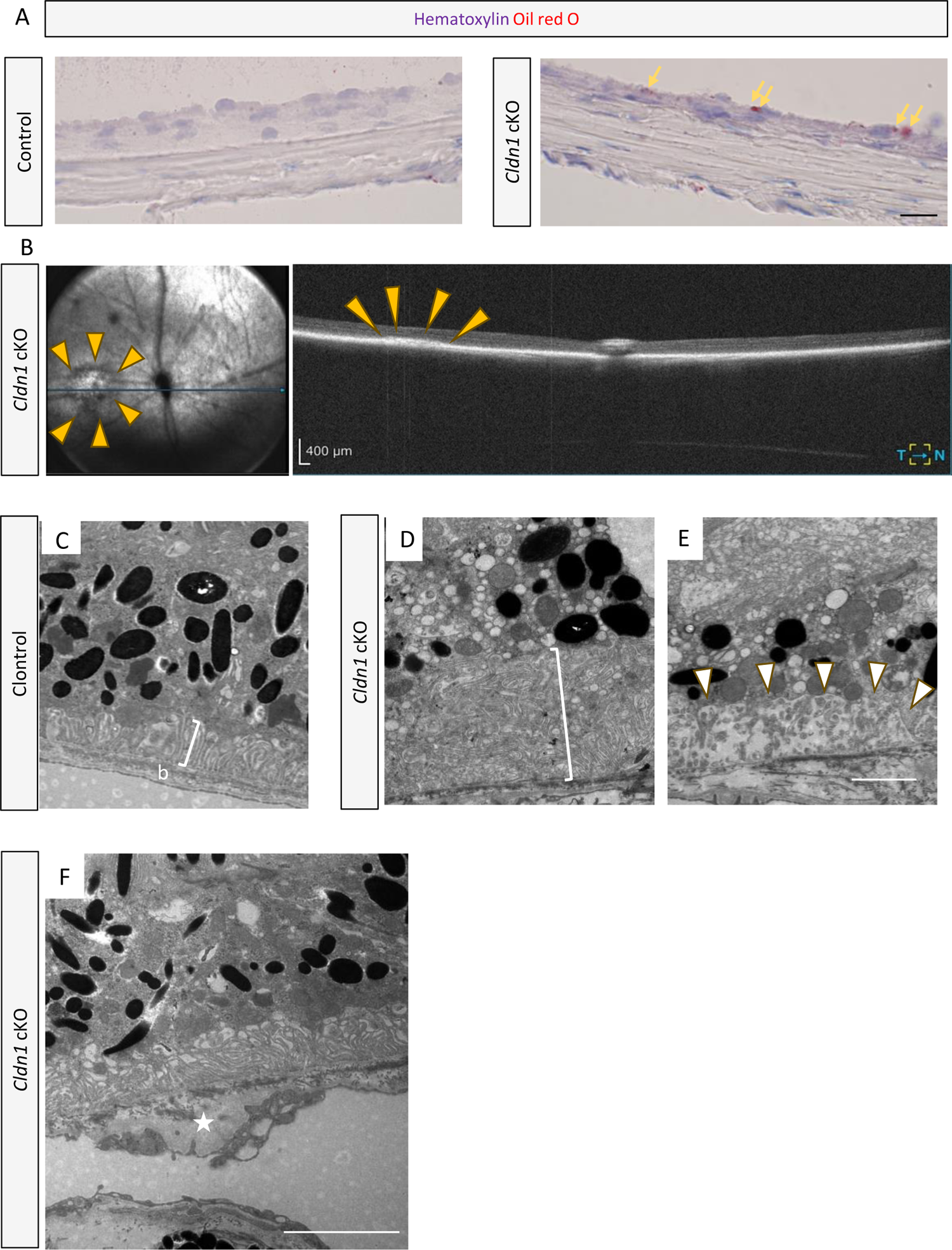
Age-dependent accumulation of lipids, lipofuscin–like vacuoles; basal laminar deposits; and membranous debris in the Bruch’s membrane by Claudin-1 deficiency. (A) Hematoxylin and oil red o staining of control and *Cldn1* cKO 16-week-old mouse after melanin bleaching. The lipid droplets (yellow arrows) were detected in the RPE of Cldn1 cKO mice. Scale bar, 20 μm. (B) Representative scanning laser ophthalmoscope (SLO) image (left) and optical coherence tomography (OCT) image (right) of 16-week-old *cldn1* cKO mouse detect sub-RPE hyperreflective region. (C-F) TEM images of RPE and Bruch’s membrane in control and *Cldn1* cKO 16-week-old mice. (n = 6 eyes per group) (C) In control mice, basal infolding (BI, bracket) was about 1-2 μm. The Bruch’s membrane (letter b) was spread uniformly in thickness under the BI. In *Cldn1* cKO mice, there were (D) thickened BL (bracket) and (E) deposits in RPE basal lamina like basal laminar deposit (BlamD) (head arrow). (D,E) Lipofuscin–like vacuoles are present near the thickened BI and BlamD. (F) Membranous debris (star) were accumulated in Bruch’s membrane of *Cldn1* cKO mice. (C-E) Equal magnification, scale bar, 2 μm. (F) Scale bar, 4 μm.

### Ectopic nuclei were observed close to subretinal drusenoid deposits (SDD) in *Cldn1* cKO mice, and macrophages within the choriocapillaris adhered to membranous debris in the Bruch’s membrane of the mice

Ectopic nuclei (EN) were often observed within photoreceptor inner segments and outer segments (IS/OS) layers of *Cldn1* cKO mice (Figure 7A). EN was DAPI-positive and appeared to be derived from the outer granular layer (Figure 7B). EN was continuously seen as photoreceptor nuclei in the TEM images (Figure 7C). Some EN had electron-dense nuclei in the TEM images (Figure 7D). Electron-dense nuclei have been known observed in apoptotic photoreceptors (Hisatomi et al., 2003; Trichonas et al., 2010). EN was observed in close to transparent membranous material containing pigments above the RPE in light microscopy (Figure 7A), and were also observed in TEM as solid accumulations of round heterogeneous vacuoles lacking internal structure resembling discs (Figure 7C,E), which were morphologically determined to be SDD. SDD was observed in 4 of 6 cKO mice eyes. In addition, macrophages with pseudopodia and vesicles in the cytoplasm adhered to thicken the Bruch’s membrane or were observed within the Bruch’s membrane in 5 out of 6 cKO mice eyes (Figure 7F,G).

**Figure 7.**
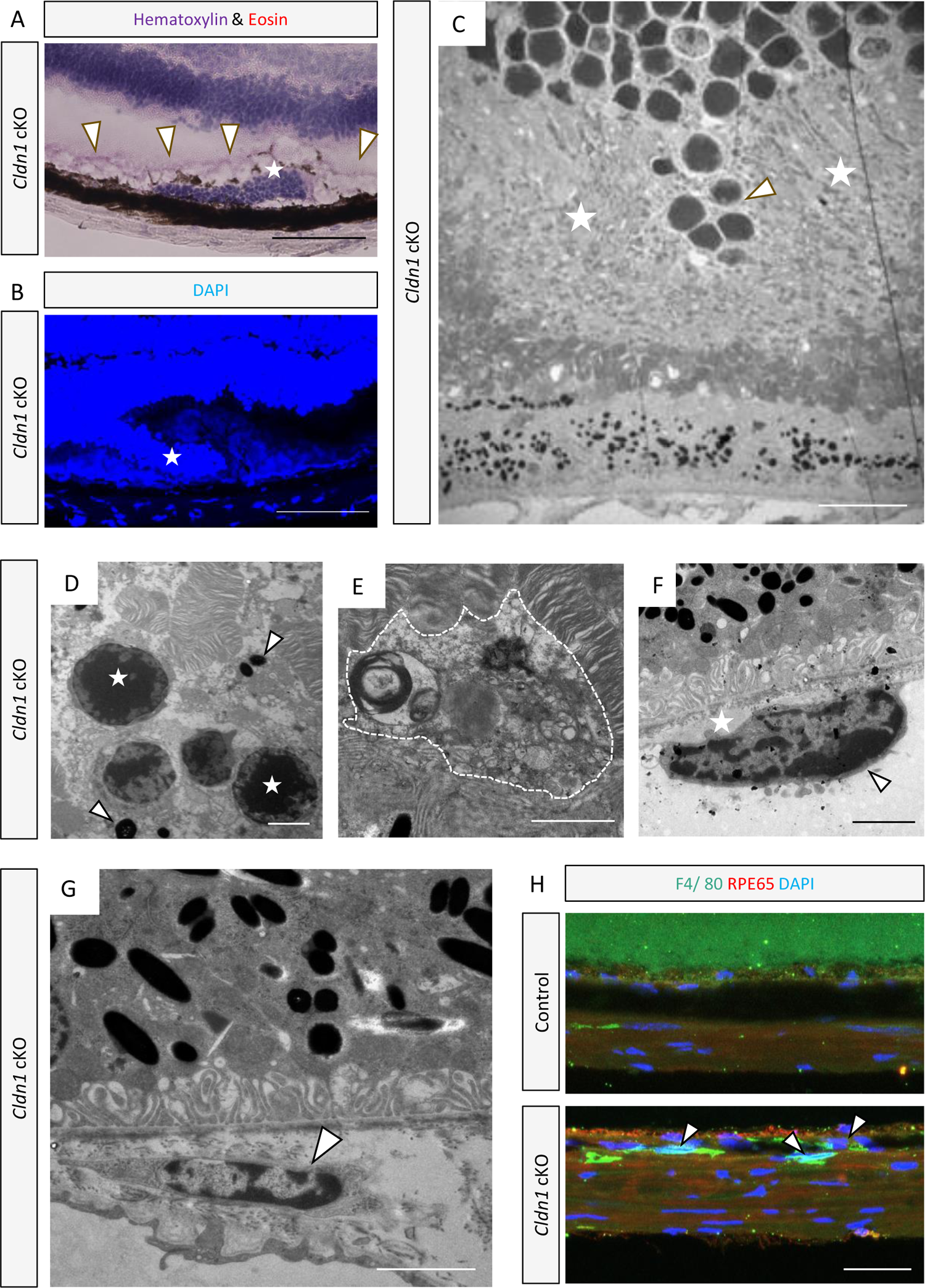
Subretinal drusenoid deposits (SDD) and ectopic nuclei (EN) as characteristic phenotypes in cldn1 cKO mice. (A) Representative hematoxylin and eosin staining and (B)DAPI staining of *Cldn1* cKO 20-week-old mouse. Scale bar, 100 μm. The stainings indicated ectopic nuclei (EN) in inner segments/outer segments (IS/OS) (star). The EN was surrounded by a membranous structure containing pigments (head arrows). (C-G) TEM images of outer layer of retina in *Cldn1* cKO 16-week-old mice. (n = 6 eyes per control and *Cldn1* cKO mice group) (C) Subretinal drusenoid deposits (SDD) (stars) was observed in the *Cldn1* cKO mice sections. SDD was found as solid materials above RPE. Loss of IS and OS was observed around the SDD, and the photoreceptor nuclei (head arrow) were disorganized and depressed toward the RPE. (D) Magnified image of EN. Some photoreceptor cell nuclei (sters) showed high electron density. RPE organelles (head arrows) were sometimes observed around EN. (E) Magnified image of another SDD. SDD (dotted line) sometimes formed between RPE and OS containing irregular round vacuoles. (F) Macrophages (head arrow) were present in Choriocapillaris. The Macrophages adhered to membranous debris of Bruch’s membrane (star). (G) The macrophage (head arrow) was observed within thicken inner collagenous layer of Bruch’s membrane. (H) 20-week-old control and *Cldn1* cKO mouse retina/choroid/sclera cross sections were stained with F4/80 (green), RPE65 (red) and DAPI (blue). Macrophages (head arrows) were present under the Bruch’s membrane and within the choriocapillaris. (C) Scale bar, 10 μm. (D-G) Scale bar, 2 μm. (H) Scale bar, 20 μm. WT; Wild type.

Immunohistochemical staining also showed F4/80-positive cells directly under the Bruch’s membrane and within the choriocapillaris (Figure 7H). There was no difference in oBRB leakage between *Cldn1* cKO and control mice (Figure S1A), while there was pooling of fluorescent dyes in SDD of *Cldn1* cKO mice (Figure S1B). In *Cldn1* cKO mice, EN, a result of photoreceptor disruption, was observed in close proximity to SDD. In addition, macrophages resided near the membranous debris in the Bruch’s membrane. The series of lipid-like deposits may induce inflammation as well as drusen containing numerous mediators of inflammation in AMD.

## Discussion

We revealed that claudin-1 is expressed in TJs of mouse RPE cells. The PCR showed that *Cldn1* was most abundantly expressed and *Cldn16* was also highly expressed in the isolated mouse RPE. There is no report of claudin-16 acting in the retina, however, claudin-16 and claudin-19 have a strong interaction and their oligomerization occurs before assembly of either claudin into the TJ in the kidney epithelial cells (Hou et al. 2009). It needs more studies in that the expression and interaction of claudin-16 in the RPE.

Claudin-1 is a transmembrane protein responsible for intercellular junctions in epithelial cells (Furuse et al., 1998) and acts as a physical barrier on the skin (Furuse et al., 2002) and intestinal (Garcia-Hernandez, Quiros and Nusrat, 2017; Suzuki, 2020) epithelial cells. The expression of the claudin family members is various in species and organs. In the human oBRB, claudin-19 plays a role in regulating the paracellular pathway (Peng et al., 2016; Wang et al., 2019). It has been suggested that oBRB breakdown is involved in the formation of retinal edema and vision loss in retinal diseases such as AMD (Bernardes et al., 2011; Joussen et al., 2021), CSC (Eandi et al., 2005; Naylor et al., 2020), and DR (Xu and Le, 2011; Omri et al., 2013; Joussen et al., 2021). However, their detailed mechanisms are still poorly understood. Based on these previous reports, we expected that claudin-1 deficiency in the mouse oBRB would cause breakdown of oBRB and leakage of serous fluid from choroid to the neural retina. However, against our expectations, *Cldn1* cKO mice did not show retinal edema, e.g. serous retinal detachment, retinal pigment epithelial detachment, or cystoid macular edema. Furthermore, there was no increase in oBRB leakage. In contrast, the cell polarity was certainly disturbed, and the RPE showed diversification of cell size, hypertrophy, and reduplication. Other claudin family members may act in a compensatory way with the loss of claudin-1 in the RPE. Thus, further studies are needed to investigate the relationship between claudin-1 and other claudin family members regarding the oBRB maintenance. In particular, more investigation of claudin family expression in young *Cldn1* cKO mice and aged *Cldn1* cKO mice using PCR and Westan blotting is needed.

In claudin-1 deficient mice, lipid accumulation (such as SDD, lipid droplets in the RPE, BlamD, and membranous debris in the Bruch’s membrane) was observed. We also observed pigment abnormalities such as RPE hypertrophy, multiple layer RPE and extracellular migration of melanin. Although age-related abnormalities of lipid metabolism and pigment abnormalities have been known as precursors and/or promoters of late AMD progression (Wang et al., 2003; Ferris et al., 2013), the notion that claudin-1 deficiency can cause those changes needs more studies. Recently, claudin has been suggested to have roles in non-junctional mechanisms. Claudin-1 promotes epithelial–mesenchymal transition (EMT) in the liver (Roehlen et al., 2022) and claudin-19 is related to degradation of internalized photoreceptor OS in the stem cell-derived RPE (Liu et al., 2021). Our study provided new insights into the possible association of claudin-1 with abnormalities in lipid metabolism and cellular senescence in the RPE.

*CX3CR1*-deficient mice have been reported to show prolonged presence of microglial cells (MC) in the subretinal space suggesting excessive photoreceptor OS phagocytosis by subretinal MC which subsequently accumulated SDD (Chuang et al., 2022). In the current study, SDD close to EN was characteristically observed in *Cldn1* deficient mice. In humans, it has been reported that a disfunction of the rod visual cycle is associated with SDD (Curcio et al., 2013). SDD is also known to be observed with photoreceptor OS and IS shortening and displacement of photoreceptor nuclei in human retinal atrophy (Chen et al., 2020). These descriptions in the previous literatures suggested that SDD in *Cldn1* cKO mice may be formed as a result of the decrease in the digestive capacity of the photoreceptor OS in the RPE. In addition, it is possible that the pathway between degradation of OS and driving cholesterol and fatty acids, which are components of the OS, to the choriocapillariswas disturbed. As a result, we were expected that AMD like extracellular deposits are accumulated.

Lipid deposition in the Bruch’s membrane impairs the diffusion of vascular endothelial growth factor A (VEGF-A) from the RPE in AMD (Holz et al., 1994). Reductions in VEGF-A levels can cause loss of choriocapillaris, finally leading to thinning of the retina and choroid (Kurihara et al., 2010, 2016; Biesemeier et al., 2014; Sohn et al., 2019). The presence of SDD was also associated with reductions in retinal thickness (Chiang et al., 2020), decreases in choroidal thickness (Switzer et al., 2012; Hogg et al., 2014; Spaide, 2018a) and risks of GA and macular neovascularization (Hogg et al., 2014; Spaide, 2018b). Reductions in retinal and choroidal thickness in adult *Cldn1* cKO mice might be related to poor diffusion of physiological mediators from the RPE. As growth factors such as VEGF-A and insulin-like growth factor 1 play crucial roles during the retinal development (Chen and Smith, 2007), 4-week-old *Cldn1* cKO mice presumably showed immature choriocapillaris and poor nutrient supplies to the retina, resulting in reductions in retinal thickness. We assumed that the growth might catch up with the adolescent 8-week-old *Cldn1* cKO mice, however; the maintenance of the retina and choroid might become difficult in mature 16-week-old *Cldn1* cKO mice due to RPE dysfunctions leading to reductions in retinal and choroidal thickness. In particular 8-week-old mice, the enlargement and diversification of RPE cells preceded the thinning of the retina and choroid, which was exactly intermediate AMD phenotype.

Inflammatory mediators such as immune complexes, complement factors, major histocompatibility complex (MHC) contained in drusen are involved in the onset of AMD (Mullins et al., 2000; Hageman et al., 2001). Mononuclear phagocytes such as monocytes and macrophages do not exist in the outer nuclear layer, sub retina, and RPE under normal conditions (Guillonneau et al., 2017). However, macrophages can be observed in the sub retina and sub RPE of GA (Espinosa-Heidmann et al., 2006; Sennlaub et al., 2013; Eandi et al., 2016), neovascular AMD (Nakashizuka et al., 2008; Yang et al., 2016), and drusen (Eandi et al., 2016). In this study, since macrophages were concentrated near the membranous debris of the Bruch’s membrane, the debris may contain inflammatory mediators like that in drusen.

## Conclusions

In summary, claudin-1 was most abundantly expressed among the claudin family in TJs of mouse RPE. RPE-specific *Cldn1* deficient mice showed early and intermediate AMD features with RPE hyperpigmentation and lipid-related metabolic abnormalities. These results indicated that members of TJ molecule claudins may play an important role on RPE homeostasis in aging.

## Acknowledgements

The authors thank Ayako Kawabata and Ryoko Shiojiri for providing animal care and genotyping and Shigeyoshi Itohara for sharing *B6-Tg(CAG-FLPe)36* mice maintained at the RIKEN BioResource Center (RBRC01834). We thank Drs. Toshihiro Nagai, Nobuko Moritoki and Tomoko Shindo at the Electron Microscope Laboratory in Keio University School of Medicine for their kind help. This work was supported by Grants-in-Aid for Scientific Research (KAKENHI, number 18 K09424) from the Ministry of Education, Culture, Sports, Science and Technology (MEXT) to Toshihide Kurihara.

---

**Supplementary Table S1.**
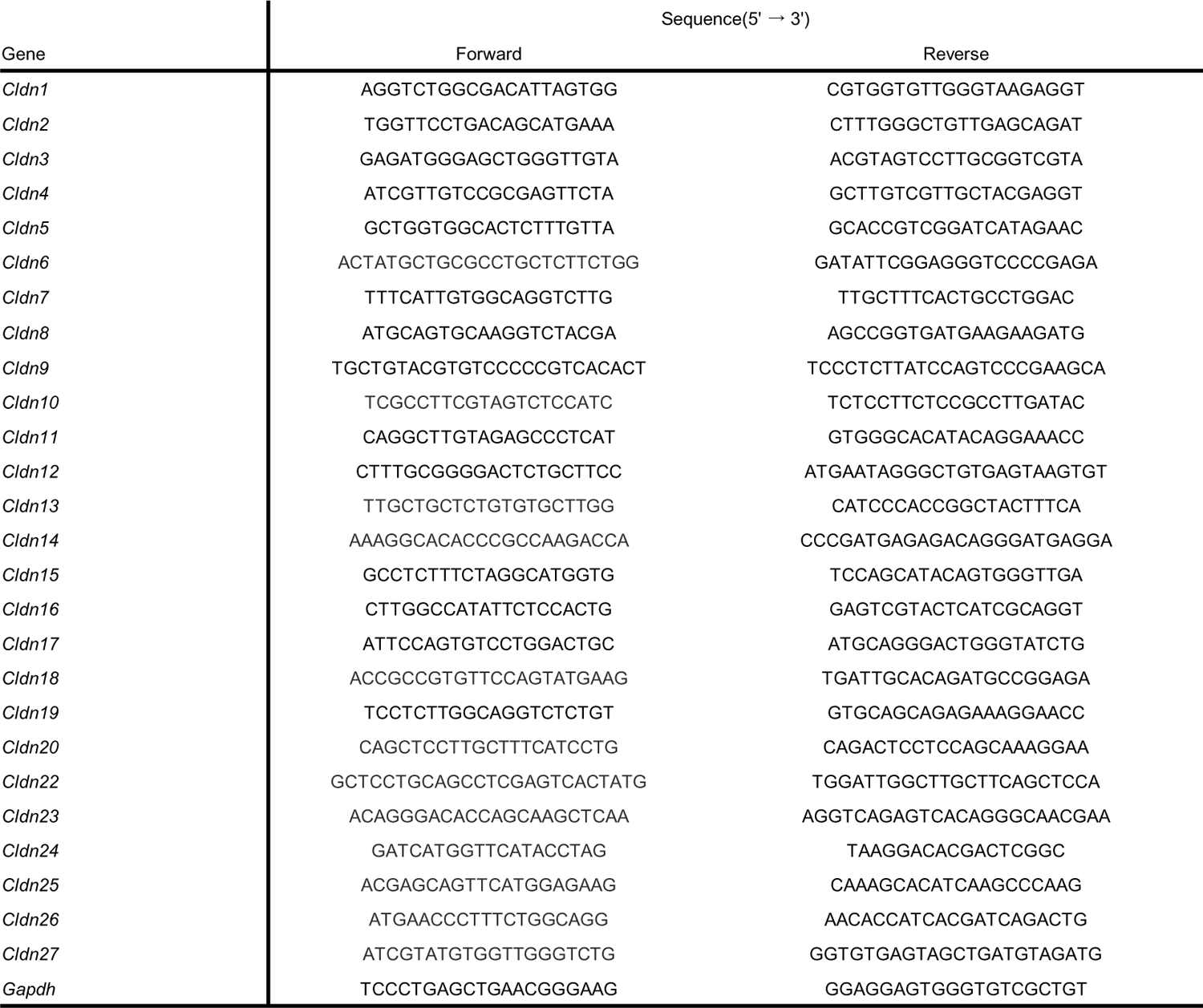
Primers Used in Quantitative PCR Analysis.

**Supplementary Figure S2.**
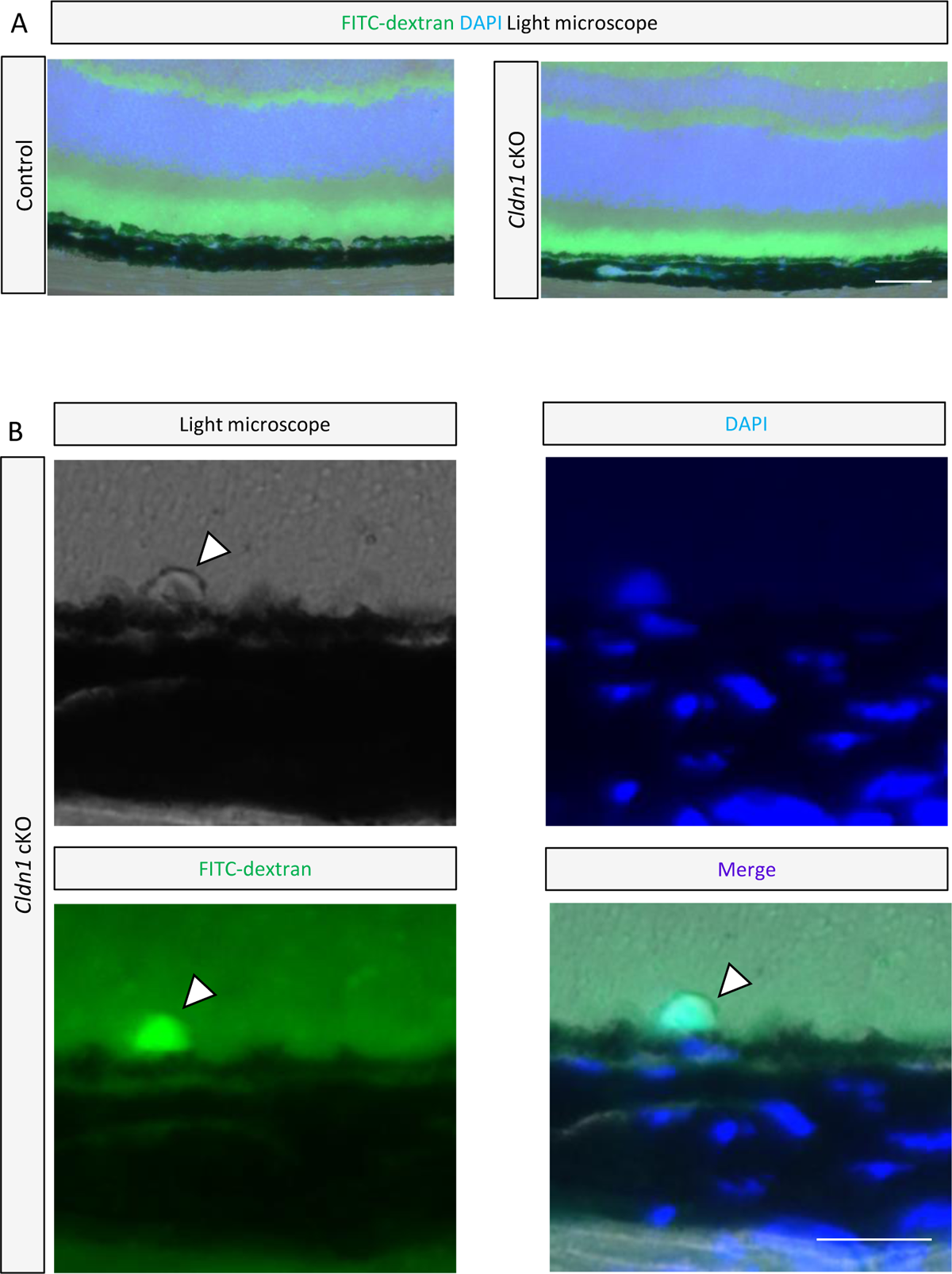
Analysis of oBRB Leakage with FITC-Dextran. (A,B) Fluorescent microscopic images of retinal sections stained with DAPI (blue) from 16-week-old control and *Cldn1* cKO mice. (A) Representative images showed no difference in FITC-dextran (green) leakage above the RPE between control and *Cldn1* cKO mice. Scale bar, 50 μm. (B) There was FITC pooling in the SDD (head arrow) of *Cldn1* cKO mice. Scale bar, 20 μm.

**Supplementary Figure S3.**
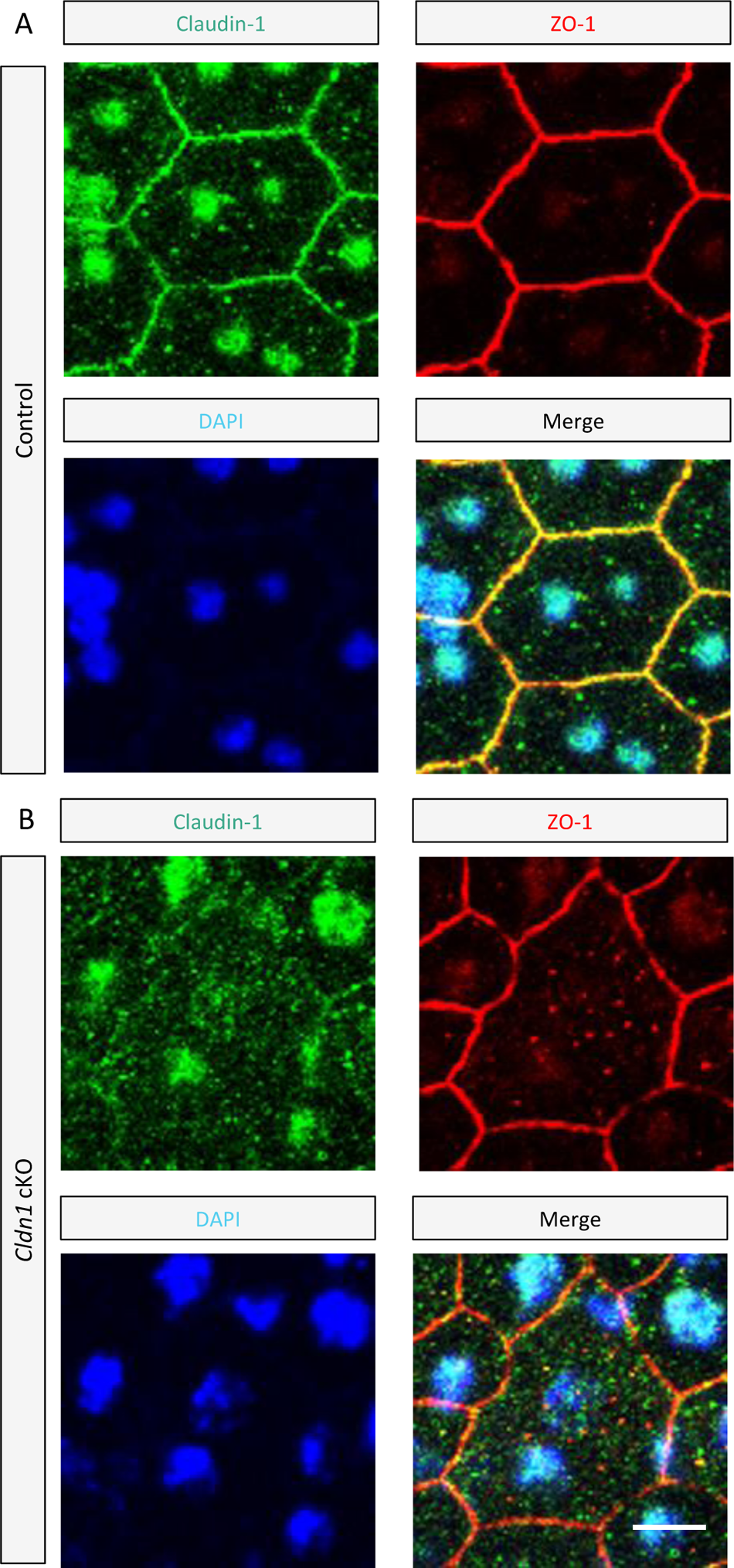
High magnification RPE structure. (A, B) RPE flat-mounts from 8-week-old control and *Cldn1* cKO mouse were stained with claudin-1 (green), ZO-1 (red) and DAPI (blue) taken at high magnification by confocal microscopy. Scale bar, 5μm.

**Supplementary Figure S4.**
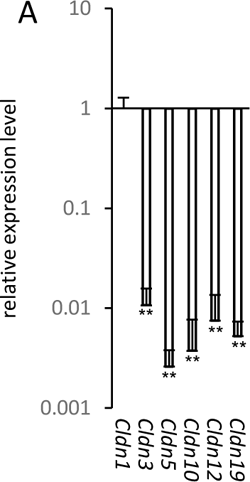
, Screening test of claudin mRNA expression in the RPE of wild type mice (C57BL/6). (A) The mRNA expression levels of claudins in RPE isolated from 12-week-old Wild type mice (± SD, n=3, 1sample 4eyes). Student t test, vs *Cldn1*, **P <0.01. The claudins has been confirmed expression in the retina from mouse cell atlas (https://bis.zju.edu.cn/MCA/index.html).

## Notes

### Competing Interest Statement

The authors have declared no competing interest.

## References

Atsugi T, et al. (2019) ‘Holocrine Secretion Occurs outside the Tight Junction Barrier in Multicellular Glands: Lessons from Claudin-1-Deficient Mice’, Journal of Investigative Dermatolog, 140(2), pp. 298–308.e5. Available at: https://www.sciencedirect.com/science/article/pii/S0022202X19331392.

Bernardes, R. et al. (2011) ‘Noninvasive Evaluation of Retinal Leakage Using Optical Coherence Tomography’, Ophthalmologica, 226(2), pp. 29–36. Available at: 10.1159/000326268.

Biesemeier, A. et al. (2014) ‘Choriocapillaris breakdown precedes retinal degeneration in age-related macular degeneration’, Neurobiology of Aging, 35(11), pp. 2562–2573. Available at: 10.1016/j.neurobiolaging.2014.05.003.

Chen, J. and Smith, L.E.H. (2007) ‘Retinopathy of prematurity’, Angiogenesis, 10(2), pp. 133–140. Available at: 10.1007/s10456-007-9066-0.

Chen, L. et al. (2020) ‘Subretinal drusenoid deposit in age-related macular degeneration: histologic insights into initiation, progression to atrophy, and imaging’, Retina (Philadelphia, Pa.), 40(4), pp. 618–631. Available at: 10.1097/IAE.0000000000002657.

Chen, M. et al. (2016) ‘Retinal pigment epithelial cell multinucleation in the aging eye – a mechanism to repair damage and maintain homoeostasis’, Aging Cell, 15(3), pp. 436–445. Available at: 10.1111/acel.12447.

Chiang, T.T.-K. et al. (2020) ‘Macular Thickness in Intermediate Age-Related Macular Degeneration Is Influenced by Disease Severity and Subretinal Drusenoid Deposit Presence’, Investigative Ophthalmology & Visual Science, 61(6), p. 59. Available at: 10.1167/iovs.61.6.59.

Chuang, J.-Z. et al. (2022) ‘Retinal pigment epithelium-specific CLIC4 mutant is a mouse model of dry age-related macular degeneration’, Nature Communications, 13(1), p. 374. Available at: 10.1038/s41467-021-27935-9.

Coşkun, M. et al. (2015) ‘Changes in the cornea related to sickle cell disease: a pilot investigation’, European Journal of Ophthalmology, 25(6), pp. 463–467. Available at: 10.5301/ejo.5000598.

Cunha-Vaz, J.G. (1976) ‘The blood-retinal barriers’, Documenta Ophthalmologica. Advances in Ophthalmology, 41(2), pp. 287–327. Available at: 10.1007/BF00146764.

Curcio, C.A. et al. (2005) ‘Esterified and unesterified cholesterol in drusen and basal deposits of eyes with age-related maculopathy’, Experimental Eye Research, 81(6), pp. 731–741. Available at: 10.1016/j.exer.2005.04.012.

Curcio, C.A. et al. (2013) ‘Subretinal Drusenoid Deposits In Non-Neovascular Age-Related Macular Degeneration: Morphology, Prevalence, Topography, And Biogenesis Model’, Retina (Philadelphia, Pa.), 33(2), p. Available at: 10.1097/IAE.0b013e31827e25e0.

Curcio, C.A. (2018) ‘Soft Drusen in Age-Related Macular Degeneration: Biology and Targeting Via the Oil Spill Strategies’, Investigative Ophthalmology & Visual Science, 59(4), pp. AMD160–AMD181. Available at: 10.1167/iovs.18-24882.

Eandi, C.M. et al. (2005) ‘ACUTE CENTRAL SEROUS CHORIORETINOPATHY AND FUNDUS AUTOFLUORESCENCE’, RETINA, 25(8), p. 989.

Eandi, C.M. et al. (2016) ‘Subretinal mononuclear phagocytes induce cone segment loss via IL-1β’, eLife. Edited by J. Nathans, 5, p. e16490. Available at: 10.7554/eLife.16490.

Espinosa-Heidmann, D.G. et al. (2006) ‘Cigarette Smoke–Related Oxidants and the Development of Sub-RPE Deposits in an Experimental Animal Model of Dry AMD’, Investigative Ophthalmology & Visual Science, 47(2), pp. 729–737. Available at: 10.1167/iovs.05-0719.

Fernandez-Godino, R., Garland, D.L. and Pierce, E.A. (2016) ‘Isolation, culture and characterization of primary mouse RPE cells’, Nature Protocols, 11(7), pp. 1206–1218. Available at: 10.1038/nprot.2016.065.

Ferris, F.L. et al. (2013) ‘Clinical Classification of Age-related Macular Degeneration’, Ophthalmology, 120(4), pp. 844–851. Available at: 10.1016/j.ophtha.2012.10.036.

Furuse, M. et al. (1998) ‘Claudin-1 and −2: Novel Integral Membrane Proteins Localizing at Tight Junctions with No Sequence Similarity to Occludin’, Journal of Cell Biology, 141(7), pp. 1539–1550. Available at: 10.1083/jcb.141.7.1539.

Furuse, M. et al. (2002) ‘Claudin-based tight junctions are crucial for the mammalian epidermal barrier: a lesson from claudin-1–deficient mice’, Journal of Cell Biology, 156(6), pp. 1099– 1111. Available at: 10.1083/jcb.200110122.

Gambril, J.A. et al. (2019) ‘Quantifying Retinal Pigment Epithelium Dysmorphia and Loss of Histologic Autofluorescence in Age-Related Macular Degeneration’, Investigative Ophthalmology & Visual Science, 60(7), pp. 2481–2493. Available at: 10.1167/iovs.19-26949.

Garcia-Hernandez, V., Quiros, M. and Nusrat, A. (2017) ‘Intestinal epithelial claudins: expression and regulation in homeostasis and inflammation’, Annals of the New York Academy of Sciences, 1397(1), pp. 66–79. Available at: 10.1111/nyas.13360.

Guillonneau, X. et al. (2017) ‘On phagocytes and macular degeneration’, Progress in Retinal and Eye Research, 61, pp. 98–128. Available at: 10.1016/j.preteyeres.2017.06.002.

Günzel, D. and Yu, A.S.L. (2013) ‘Claudins and the Modulation of Tight Junction Permeability’, Physiological Reviews, 93(2), pp. 525–569. Available at: 10.1152/physrev.00019.2012.

Hageman, G.S. et al. (2001) ‘An Integrated Hypothesis That Considers Drusen as Biomarkers of Immune-Mediated Processes at the RPE-Bruch’s Membrane Interface in Aging and Age-Related Macular Degeneration’, Progress in Retinal and Eye Research, 20(6), pp. 705–732. Available at: 10.1016/S1350-9462(01)00010-6.

Haimovici, R. et al. (2001) ‘The Lipid Composition of Drusen, Bruch’s Membrane, and Sclera by Hot Stage Polarizing Light Microscopy’, Investigative Ophthalmology & Visual Science, 42(7), pp. 1592–1599.

Hisatomi, T. et al. (2003) ‘Clearance of Apoptotic Photoreceptors: Elimination of Apoptotic Debris into the Subretinal Space and Macrophage-Mediated Phagocytosis via Phosphatidylserine Receptor and Integrin αvβ3’, The American Journal of Pathology, 162(6), pp. 1869–1879. Available at: 10.1016/S0002-9440(10)64321-0.

Hou, J. et al. (2009) ‘Claudin-16 and claudin-19 interaction is required for their assembly into tight junctions and for renal reabsorption of magnesium’, Proceedings of the National Academy of Sciences,106(36),pp.15350–15355.Available at: https://www.pnas.org/doi/10.1073/pnas.0907724106

Niwa, H. et al. (1991) ‘Efficient selection for high-expression transfectants with a novel eukaryotic vector’, Gene, 108(2), pp. 193–199. Available at: 10.1016/0378-1119(91)90434-D.

Hogg, R.E. et al. (2014) ‘Clinical Characteristics of Reticular Pseudodrusen in the Fellow Eye of Patients with Unilateral Neovascular Age-Related Macular Degeneration’, Ophthalmology, 121(9), pp. 1748–1755. Available at: 10.1016/j.ophtha.2014.03.015.

Holz, F.G. et al. (1994) ‘Analysis of Lipid Deposits Extracted From Human Macular and Peripheral Bruch’s Membrane’, Archives of Ophthalmology, 112(3), pp. 402–406. Available at: 10.1001/archopht.1994.01090150132035.

Iacovelli, J. et al. (2011) ‘Generation of Cre Transgenic Mice with Postnatal RPE-Specific Ocular Expression’, Investigative Ophthalmology & Visual Science, 52(3), pp. 1378–1383. Available at: 10.1167/iovs.10-6347.

Joussen, A.M. et al. (2021) ‘Angiopoietin/Tie2 signalling and its role in retinal and choroidal vascular diseases: a review of preclinical data’, Eye, 35(5), pp. 1305–1316. Available at: 10.1038/s41433-020-01377-x.

Kaur, C., Foulds, W.S. and Ling, E.A. (2008) ‘Blood–retinal barrier in hypoxic ischaemic conditions: Basic concepts, clinical features and management’, Progress in Retinal and Eye Research, 27(6), pp. 622–647. Available at: 10.1016/j.preteyeres.2008.09.003.

Kurihara, T. et al. (2010) ‘von Hippel-Lindau protein regulates transition from the fetal to the adult circulatory system in retina’, Development, 137(9), pp. 1563–1571. Available at: 10.1242/dev.049015.

Kurihara, T. et al. (2016) ‘Hypoxia-induced metabolic stress in retinal pigment epithelial cells is sufficient to induce photoreceptor degeneration’, eLife. Edited by J. Nathans, 5, p. e14319. Available at: 10.7554/eLife.14319.

Li, S. et al. (2011) ‘Retro-orbital injection of FITC-dextran is an effective and economical method for observing mouse retinal vessels’, Molecular Vision, 17, pp. 3566–3573.

Liu, F. et al. (2021) ‘Knockdown of Claudin-19 in the Retinal Pigment Epithelium Is Accompanied by Slowed Phagocytosis and Increased Expression of SQSTM1’, Investigative Ophthalmology & Visual Science, 62(2), p. 14. Available at: 10.1167/iovs.62.2.14.

Matsuda, M., Yee, R.W. and Edelhauser, H.F. (1985) ‘Comparison of the corneal endothelium in an American and a Japanese population’, Archives of Ophthalmology (Chicago, Ill.: 1960), 103(1), pp. 68–70. Available at: 10.1001/archopht.1985.01050010072023.

Mineta, K. et al. (2011) ‘Predicted expansion of the claudin multigene family’, FEBS Letters, 585(4), pp. 606–612. Available at: 10.1016/j.febslet.2011.01.028.

Mitchell, P. et al. (2018) ‘Age-related macular degeneration’, The Lancet, 392(10153), pp. 1147–1159. Available at: 10.1016/S0140-6736(18)31550-2.

Mullins, R.F. et al. (2000) ‘Drusen associated with aging and age-related macular degeneration contain proteins common to extracellular deposits associated with atherosclerosis, elastosis, amyloidosis, and dense deposit disease’, The FASEB Journal, 14(7), pp. 835–846. Available at: 10.1096/fasebj.14.7.835.

Nakashizuka, H. et al. (2008) ‘Clinicopathologic Findings in Polypoidal Choroidal Vasculopathy’, Investigative Ophthalmology & Visual Science, 49(11), pp. 4729–4737. Available at: 10.1167/iovs.08-2134.

Naylor, A. et al. (2020) ‘Tight Junctions of the Outer Blood Retina Barrier’, International Journal of Molecular Sciences, 21(1), p. 211. Available at: 10.3390/ijms21010211.

Omri, S. et al. (2013) ‘PKCζ Mediates Breakdown of Outer Blood-Retinal Barriers in Diabetic Retinopathy’, PLOS ONE, 8(11), p. e81600. Available at: 10.1371/journal.pone.0081600.

Ortolan, D. et al. (2022) ‘Single-cell–resolution map of human retinal pigment epithelium helps discover subpopulations with differential disease sensitivity’, Proceedings of the National Academy of Sciences, 119(19), p. e2117553119. Available at: 10.1073/pnas.2117553119.

Peng, S. et al. (2011) ‘Claudin-19 and the Barrier Properties of the Human Retinal Pigment Epithelium’, Investigative Ophthalmology & Visual Science, 52(3), pp. 1392–1403. Available at: https://iovs.arvojournals.org/article.aspx?articleid=2126564

Peng, S. et al. (2016) ‘Claudin-3 and claudin-19 partially restore native phenotype to ARPE-19 cells via effects on tight junctions and gene expression’, Experimental Eye Research, 151, pp. 179–189. Available at: 10.1016/j.exer.2016.08.021.

Roehlen, N. et al. (2022) ‘A monoclonal antibody targeting nonjunctional claudin-1 inhibits fibrosis in patient-derived models by modulating cell plasticity’, Science Translational Medicine, 14(676), p. eabj4221. Available at: 10.1126/scitranslmed.abj4221.

Runkle, E.A. and Antonetti, D.A. (2011) ‘The Blood-Retinal Barrier: Structure and Functional Significance’, in S. Nag (ed.) The Blood-Brain and Other Neural Barriers: Reviews and Protocols. Totowa, NJ: Humana Press (Methods in Molecular Biology), pp. 133–148. Available at: 10.1007/978-1-60761-938-3_5.

Sakaguchi, H. et al. (2002) ‘Clusterin is Present in Drusen in Age-related Macular Degeneration’, Experimental Eye Research, 74(4), pp. 547–549. Available at: 10.1006/exer.2002.1186.

van der Schaft, T.L. et al. (1993) ‘Basal laminar deposit in the aging peripheral human retina’, Graefe’s Archive for Clinical and Experimental Ophthalmology, 231(8), pp. 470–475. Available at: 10.1007/BF02044234.

Sennlaub, F. et al. (2013) ‘CCR2+ monocytes infiltrate atrophic lesions in age-related macular disease and mediate photoreceptor degeneration in experimental subretinal inflammation in Cx3cr1 deficient mice’, EMBO Molecular Medicine, 5(11), pp. 1775–1793. Available at: 10.1002/emmm.201302692.

Sohn, E.H. et al. (2019) ‘Choriocapillaris Degeneration in Geographic Atrophy’, The American Journal of Pathology, 189(7), pp. 1473–1480. Available at: 10.1016/j.ajpath.2019.04.005.

Spaide, R.F. (2018a) ‘DISEASE EXPRESSION IN NONEXUDATIVE AGE-RELATED MACULAR DEGENERATION VARIES WITH CHOROIDAL THICKNESS’, RETINA, 38(4), p. 708. Available at: 10.1097/IAE.0000000000001689.

Spaide, R.F. (2018b) ‘IMPROVING THE AGE-RELATED MACULAR DEGENERATION CONSTRUCT: A New Classification System’, RETINA, 38(5), p. 891. Available at: 10.1097/IAE.0000000000001732.

Suzuki, T. (2020) ‘Regulation of the intestinal barrier by nutrients: The role of tight junctions’, Animal Science Journal, 91(1), p. e13357. Available at: 10.1111/asj.13357.

Switzer, D.W. et al. (2012) ‘Segregation of ophthalmoscopic characteristics according to choroidal thickness in patients with early age-related macular degeneration’, Retina (Philadelphia, Pa.), 32(7), pp. 1265–1271. Available at: 10.1097/IAE.0b013e31824453ac.

Trichonas, G. et al. (2010) ‘Receptor interacting protein kinases mediate retinal detachment-induced photoreceptor necrosis and compensate for inhibition of apoptosis’, Proceedings of the National Academy of Sciences, 107(50), pp. 21695–21700. Available at: 10.1073/pnas.1009179107.

Watzke, R.C. et al. (1993) ‘Morphometric analysis of human retinal pigment epithelium: correlation with age and location’, Current Eye Research, 12(2), pp. 133–142. Available at: 10.3109/02713689308999481

Wang, J.J. et al. (2003) ‘Risk of Age-Related Macular Degeneration in Eyes With Macular Drusen or Hyperpigmentation: The Blue Mountains Eye Study Cohort’, Archives of Ophthalmology, 121(5), pp. 658–663. Available at: 10.1001/archopht.121.5.658.

Wang, S.-B. et al. (2019) ‘Disease-associated mutations of claudin-19 disrupt retinal neurogenesis and visual function’, Communications Biology, 2(1), p. 113. Available at: 10.1038/s42003-019-0355-0.

Wu, J. et al. (2006) ‘Identification of new claudin family members by a novel PSI-BLAST based approach with enhanced specificity’, Proteins: Structure, Function, and Bioinformatics, 65(4), pp. 808–815. Available at: 10.1002/prot.21218.

Xu, H.-Z. and Le, Y.-Z. (2011) ‘Significance of Outer Blood–Retina Barrier Breakdown in Diabetes and Ischemia’, Investigative Ophthalmology & Visual Science, 52(5), pp. 2160–2164. Available at: 10.1167/iovs.10-6518.

Yang, Y. et al. (2016) ‘Macrophage polarization in experimental and clinical choroidal neovascularization’, Scientific Reports, 6(1), p. 30933. Available at: 10.1038/srep30933.

Yardeni, T. et al. (2011) ‘Retro-orbital injections in mice’, Lab animal, 40(5), pp. 155–160. Available at: 10.1038/laban0511-155.

Zhang, C. et al. (2019) ‘Erythropoietin protects outer blood-retinal barrier in experimental diabetic retinopathy by up-regulating ZO-1 and occludin’, Clinical & Experimental Ophthalmology, 47(9), pp. 1182–1197. Available at: 10.1111/ceo.13619.

